# Links between mouse and vole social networks and their gut microbiomes support predictions from metacommunity theory

**DOI:** 10.1101/2020.08.18.256370

**Authors:** Joël W. Jameson, Steven W. Kembel, Denis Réale

## Abstract

Metacommunity theory predicts that strongly connected individuals will harbour similar gut microbiomes (GMs) and affiliating with more individuals should increase GM diversity. Additionally, cross-species bacterial transmission may play a role in how interspecific interactions affect host community dynamics. We tracked sympatric mice (*Peromyscus maniculatus*) and voles (*Myodes gapperi*) and constructed social networks for each species and both species together. We tested whether: 1) similarity in GM composition between individuals correlates with their social proximity within and across species; 2) GM diversity correlates with a host’s number of conspecific or heterospecific neighbours. We could not differentiate associations between GM composition and mouse social proximity or habitat. In voles, social proximity explained part of the GM composition. GM composition associated with interspecific social proximity, and mouse GM diversity correlated with number of vole neighbours. Contributions of host-host bacterial transmission to the GM partly follow metacommunity theory but depend on host species.

## INTRODUCTION

There is growing interest in the ways that social interactions among animals relate to their microbiome. The gut microbiome (GM) can play an important role in digestion of carbohydrates (Neish, 2009), lipid storage (Bäckhed et al., 2004), immune activation (Karmarkar and Rock, 2013), and development of the central nervous system (Desbonnet et al., 2014; Heijtz et al., 2011; Hoban et al., 2016; Neufeld et al., 2011). All these effects have downstream consequences for social behaviour (Desbonnet et al., 2014; Gacias et al., 2016). Dysbiosis of the GM is associated with autism spectrum disorder (Mohamadkhani, 2018), depression, anxiety and social impairment (Amato, 2013; De Palma et al., 2015; Golubeva et al., 2017) as well as intestinal pathologies including Crohn’s disease (Joossens et al., 2011) and irritable bowl syndrome (Kennedy et al., 2014). Given the microbiome’s association with host health and social behaviour, understanding how GM composition relates to interactions among hosts can help us better understand population dynamics with implications for conservation.

Considering the host as a habitat patch inhabited by a community of species, the microbiome, allows the use of metacommunity theory to explain at least part of how host social interactions affect GM composition (Amato, 2013; Miller et al., 2018). Following metacommunity theory (Leibold et al., 2004; MacArthur and Wilson, 1967), if microbes disperse between hosts, then similar microbial communities should inhabit closely connected hosts, and more isolated hosts should show lower diversity. Abundant evidence demonstrates the capacity for host-host bacterial transmission to shape the GM. For example, in humans, the mother transfers perinatal GM via the uterus, vagina and milk (Amato, 2013). In adulthood, individuals that live together share more similar GMs than non-housemates (Song et al., 2013) with greater similarity and increased bacterial diversity in the GMs of close married couples (Dill-McFarland et al., 2019). Additionally, for people that live alone, higher levels of social interactions increase GM diversity (Dill-McFarland et al., 2019). Also, mice that share the same cage develop similar GM compositions, either through direct contacts or indirectly via the environment (Alexander et al., 2006; Antonopoulos et al., 2009; Hildebrand et al., 2013).

Besides studies on humans and laboratory animals, social network studies of wild animals provide further evidence for a strong role of host-host interactions in shaping the GM composition and diversity. During periods of elevated social aggregation, and regardless of diet, degree of social contacts among chimpanzees (*Pan troglodytes*) explained 7.6% of variation in GM richness and 4.8% of variation in GM composition (Moeller et al., 2016). Tung et al. (Tung et al., 2015) found that 19% of the GM of baboons (*Papio cynocephalus*) was attributable to differences among social groups. Antwis et al. (Antwis et al., 2018) identified within- and across-band variation in the GM composition of semi-feral ponies (*Equus ferus caballus*) consistent with social proximity, with band explaining 14% of GM composition. In red-bellied lemurs (*Eulemur rubriventer*) up to 28% of the variance in the GMs was attributable to social group, once controlling for time, age and sex but not diet (Raulo et al., 2018). Interestingly, Raulo et al. observed a negative correlation between individual sociality (i.e. propensity of social grooming) and GM diversity, which they claim counters metacommunity theory. However, metacommunity theory predicts greater diversity at intermediate levels of dispersal (Kunin, 1998; Mouquet et al., 2006; Venail et al., 2010) meaning GM diversity may be lower in highly social species. Strong evidence for an association between social interactions and host GM composition and diversity, thus, seems consistent with current metacommunity theory. This evidence, however, comes from studies on highly social species including humans and other primates, and domestic or laboratory animals (Alexander et al., 2006; Antwis et al., 2018; Hildebrand et al., 2013; Moeller et al., 2016; Raulo et al., 2018; Tung et al., 2015). We must extend our knowledge to other social architectures to fully understand the role of dispersal in shaping the GM.

Past studies have also focused on a single species at a time in widely varied geographic locations, but bacterial transmission may occur across co-occurring species. Adult humans share a more similar skin microbiome with their dogs than with that of other dogs, and dog owners skin microbiomes resemble each other more than those of people without dogs (Song et al., 2013). Additionally, experimental GM transfers can successfully humanize the GM of mice (Turnbaugh et al., 2009; Wrzosek et al., 2018), indicating the possibility of cross-species microbial dispersal to shape the GM. Cross-species microbial transfer may occur directly through predator-prey relationships (Moeller et al., 2017), or through competition and agonistic interactions, or indirectly through shared habitats. Agonistic interactions have been observed between several small mammal species in dyadic neutral field tests (Banks et al., 1979; Falkenberg and Clarke, 1998; Grant, 1972; Rowley and Christian, 1976; Rychlik and Zwolak, 2006). In the wild, agonistic interactions among species are suggested to facilitate coexistence (Rychlik and Zwolak, 2006). We suspect that competitive interactions between two species or their sympatric use of resources can facilitate the transmission of GM members from one species to another, which could have strong eco-evolutionary consequences. For instance, transfer of beneficial or pathogenic bacteria could change the competition game between the two species. It is also important to understand the extent of the role that some individuals might play in that interspecies transfer, as it may create fitness differences among individuals within each species. However, the extent to which such transmission occurs in the wild has not been tested.

Here we examine the role of social proximity in two sympatric and ecologically similar species of rodents, the deer mouse (*Peromyscus maniculatus*) and the red-backed vole (*Myodes gapperi*). Both species are common and widespread in North America, have similar habitat preferences, population dynamics, and reproductive strategies (Harper and Austad, 2001). Both species mate promiscuously, and females are territorial while males are not (Mihok, 1981). Consequently, their home ranges overlap considerably, enabling within- and cross-species bacterial dispersal that could affect the GM. Additionally, neither species forms strong social groups (Kawata, 1985; Millar and Derrickson, 1992), though deer mice show some plasticity in social gregariousness (Millar and Derrickson, 1992; Savidge, 1974), so testing associations between GM and social proximity in this system should afford a better understanding of the extent to which metacommunity theory can predict GM composition. Specifically, we hypothesize that 1) similarity in GM composition between individuals will be correlated with their likelihood of association, at the within- and across-species levels; and 2) an individual’s GM diversity will increase with the number of conspecific or heterospecific neighbours.

## METHODS

This study took place from 14 July—2 August 2016 (20 days) on Harbour Island in the Winnipeg River basin, Ontario (50°2.580’N, 94°40,436’W). We captured mice and voles with Longworth and BIOEcoSS (BIOEcoSS Ltd.) live traps. We placed 192 traps at 96 stations (8 × 12) distanced 10 m apart which provided a capture grid (9600 m^2^) large enough to estimate individual home range sizes (Thompson et al., 2009; Vanderwel et al., 2010; Wolff, 1985; Wood et al., 2010). Weather permitting, we conducted captures every second day. We opened the traps in the evening and checked and closed them at 6:00 a.m. Before each trap deployment, we cleaned and sterilized the traps with 70% ethanol and baited them with 4 g of carrot and 1 ml of peanut butter. For each capture, we recorded age as one of three classes (juvenile, subadult, adult) based on pelage colour (Naughton, 2012; Osgood, 1909) and sex based on urogenital distance. After processing, we returned animals to their precise trapping location.

When first captured, we outfitted each animal with a subcutaneous, passive integrated transponder (PIT) tag. This allowed us to rapidly identify recaptured animals, and to collect movement data by deploying 16 RFID data loggers at equal distances throughout the capture grid starting on Day 12 (July 25). Data loggers consisted of a ring-shaped antenna placed onto a wooden arena (20 × 20 × 5 cm). The arena contained closed compartments filled with peanut butter. The compartments were perforated so animals could smell the bait but not eat it. We regularly replaced the bait in these compartments. The floor of the arena was a plastic grid elevated 1–2 cm above the ground to prevent accumulation of feces within the arena and prevent unwanted cross-contamination among animals. When an animal passed through the antenna ring, the data logger recorded its identity and time of detection. We counted multiple consecutive detections for an individual at the same RFID station and on the same day as a single detection.

We obtained a dual-species social network based on the compiled capture and PIT tag data from both species. Since traps and data loggers do not sample equally, we randomly selected a single detection from each individual on capture days but used all the detections on non-capture days. This allowed us to maximize the data used in estimating social networks while balancing survey effort. We also only retained individuals with at least five days of data. To estimate the social network we used a novel approach described in (Wanelik and Farine, 2019) wherein we use logistic models to model sex- and species-specific space use. Using the data from both species, we first built a distance matrix based on each individuals’ centroid and, for each pair of individuals in the matrix, we estimated their probability of occurring at each of 100 distance slices between their centroids. We then took the parallel minima and maxima of those probabilities and, finally, the ratio of the sum of the parallel minima and the sum of the parallel maxima gives a measure of the encounter probability between each pair of individuals. The matrix of encounter probabilities (hereafter “social matrix”) can then be graphically represented as a social network where the edge weights correspond to the encounter probability in the matrix. We then subsampled the social matrix to obtain a social matrix for voles, and another for mice. We assessed whether the dual (i.e. mouse and vole) and single-species social networks were non-randomly structured using a Monte Carlo approach wherein we generated 1000 random networks and compared the coefficients of variance of our observed social matrices to those of the random social matrices. We transformed the social matrices into social distance matrices by taking the log of the inverse of the social matrices. We did this because some subsequent analyses required input matrices to be in this format and it made interpretation of results easier since microbiome composition was summarized as a dissimilarity (or distance) matrix. After transformation, the log(social matrix) and the distance matrix were perfectly negatively correlated (R^2^=1, see supplementary information).

To account for the potential confounding effect of shared habitat on the GM, we collected habitat data at each trapping station and included information on habitat composition in our analyses testing our first hypothesis on GM composition, and on habitat composition and diversity when testing our second hypothesis on GM diversity. At each station, within a 5 m-radius circle centered on the trap, we characterized the overstory by measuring the abundance of all tree species with a diameter at breast height (dbh)>10 cm as well as the percent total canopy cover. We characterized the understory by measuring, within a 2 m-radius circle centered on the trap, the percent cover of all other plant species. We did not identifie mosses and lichens to species. We also noted the percent cover of deadwood, tree stumps, litter, and rock. Finally, we measured soil moisture and pH at a distance of one meter from the trap in each cardinal direction. We simplified the data by removing overstory species with an abundance <5% of the total overstory data and by removing understory species with an abundance <1% of the understory data. We Hellinger-transformed the habitat community data and summarized the variance in habitat data with principal component analysis (PCA) (Borcard et al., 2011). The first three PCA components cumulatively accounted for 43% of the variance in habitat data so we used their scores (Habitat-PC1, Habitat-PC2, Habitat-PC3) as habitat variables in subsequent analyses. Finally, to obtain a single value for each score per individuals, we averaged the habitat scores over each individual’s detection events. We measured habitat diversity using Simpson’s diversity index.

We collected samples from feces left in the trap by the captured individual. We collected two to three fecal samples per animal throughout the study period. We amplified and sequenced the V4-V5 regions (~410 bp) of the 16S rRNA gene. We identified bacterial sequences to amplicon sequence variants (ASV) with DADA2 (Callahan et al., 2016) and performed taxonomic assignment with SILVA database (V128). Details on sample storage, DNA extraction, amplification and sequencing are described in (Jameson et al., 2020). Quality control procedures applied to the sequence data are described in supplementary methods.

As a general test of our first hypothesis, we used a Procrustes analysis (least-squares orthogonal mapping; function protest, package Vegan; Oksanen et al., 2019; Peres-Neto and Jackson, 2001) to test for structural similarities between the microbiome and social distance matrices. We also ran the same analysis testing microbiome dissimilarity against habitat dissimilarity (Euclidean distance). This analysis generates a measure of fit (m_12_^2^), which is the sum of squared deviations between two matrix configurations, from which it derives a correlation index (t). It then uses a permutation test to calculate the probability (p-value) of observing our m_12_^2^ (Oksanen et al., 2019; Peres-Neto and Jackson, 2001). We ran these two tests for each species’ social distance matrix and for the dual-species social distance matrix. We also ran this test on the subset of interspecific pairs from the dual-species social distance matrix which allowed us to focus the analysis on between-species interactions.

We then used a constrained redundancy analysis (RDA; function rda, package Vegan; Borcard et al., 2011) as a more thorough test of our first hypothesis. This analysis allowed us to identify gradients of microbiome composition potentially associated with social structure and to identify components of social structure correlated with the microbiome and not confounded with shared habitat. To extract gradients of variation in social distance that we could use in the RDA, we ran principal coordinate analysis (PCoA; function pcoa, package ape; Paradis et al., 2020) on social distance and used the scores of the first three axes (Social-PCo1, Social-PCo2, Social-PCo3) in the RDA analyses. These axes cumulatively represented 47%, 43%, and 16% of the variance in the mouse, vole and dual-species networks respectively. We ran separate RDA analyses for each species and for the dual-species network. Generally, we included the Hellinger-transformed microbiome community as the dependent variable, and the social and habitat scores as independent variables. We also included sex in the analysis for voles, but not for mice as the mouse data for this analysis contained a single female. Additionally, for the dual species model, we first removed the variance associated with species and sex by conditioning for these variables in a partial RDA. We used forward stepwise model selection and set the selection procedure to stop when the adjusted R^2^ reached that from the full model. We used an ANOVA-like permutation test to assess the significance of the models and of each independent variable in the models. To understand which social and habitat variables were confounded, we ran the above analyses without the habitat variables, and once more conditioning on the habitat variables (Figures 2–5).

**Figure 1.**
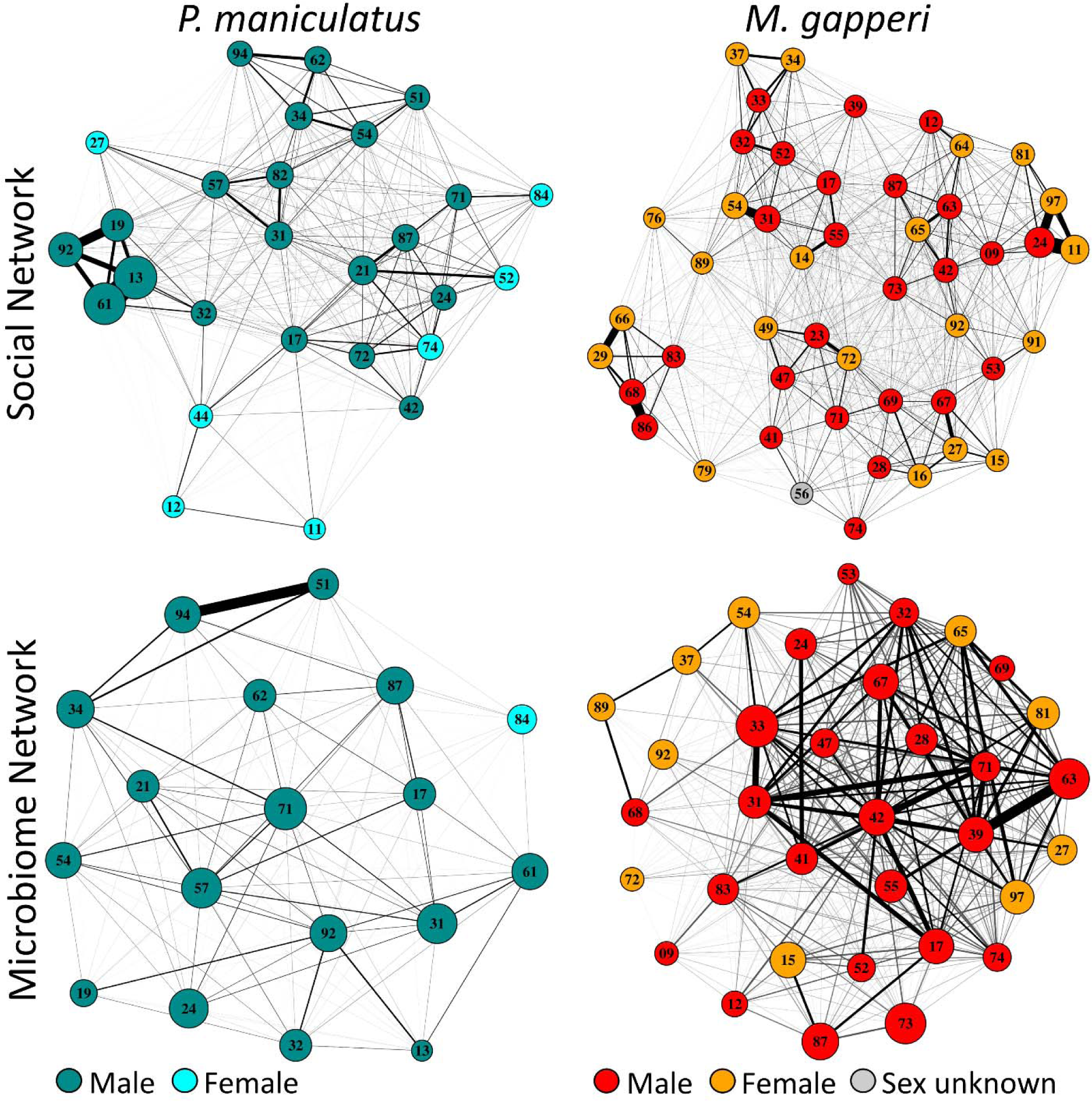
Social network (above) and microbiome distance network (below) of *P. maniculatus* and *M. gapperi*. Vertex size is proportional to degree centrality in social networks and GM Shannon’s diversity in microbiome networks. Edge width is proportional to the strength of association in social networks and to compositional similarity in GM networks. To improve visualization of the networks, edge colour is made proportional to the strength of the association. Edge weights were divided into 10% quantile categories and each category was assigned a colour following a gradient from white (weakest association) to black (strongest association). Numbers identify individuals and vertex colours identify species and sex.

**Figure 2.**
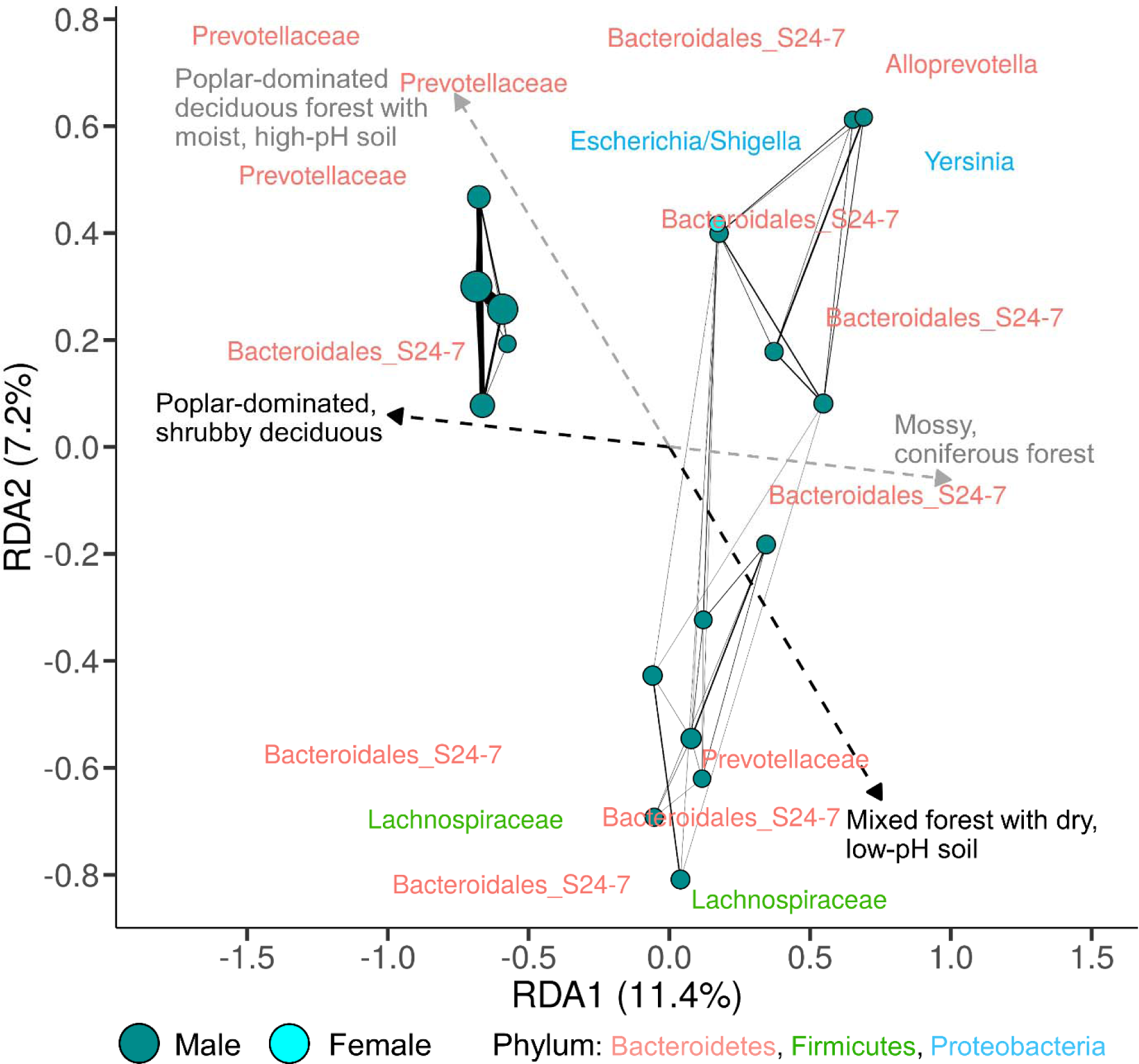
Biplot of RDA of the association between GM composition and habitat in *P. maniculatus*. The model was selected from a full model that included social network and habitat as explanatory factors. Variables correlated with GM composition and the direction and strength of the correlations are indicated by arrows. Habitat variables used were scores of a PCA of the habitat data and are represented above by decriptions of vegetation structure along these PCA axes. Social variables used were scores of a PCoA on social distance. Individuals with the strongest social associations (above the third quartile of social distances) are connected by lines, line width is proportional to the strength of association, and point size is proportional to degree centrality in the mouse social network. The 10 most strongly loaded bacterial ASVs along each axis (identified to the finest-level taxonomic annotation for that ASV) are plotted (duplicate ASVs were removed).

To address the effect of an individuals’ position in the network relative to conspecifics and heterospecifics (i.e. number of social neighbours), we obtained three measures of degree centrality for each species. We used degree centrality since it is equivalent to other network parameters in predicting information flow but more suited to potentially incomplete datasets due to its simplicity and lower sensitivity to error (Drewe and Perkins, 2015). For each individual of both species, we first measured the degree centrality of an individual in its conspecific network, and its degree centrality taken from the dual-species network. For the third measure, we obtained that individual’s degree centrality relative to the heterospecific network.

We then measured Shannon’s, Simpson’s, and richness indices of microbiome diversity (function diversity, package vegan). Simpson’s diversity is more sensitive to abundant species, richness to rare species, and Shannon’s diversity to rare and abundant species equally (Morris et al., 2014). We tested our hypothesis on GM diversity with all three indices as this allowed us to better understand which component of diversity was affected by social transmission. We then ran linear models (package stats) with microbiome diversity as the dependent variable and degree centrality, Habitat-PC1, Habitat-PC2, Habitat-PC3, and Simpson’s habitat diversity as independent variables. When necessary, we log-transformed microbiome diversity to meet normality assumptions. We ran separate models for each measure of microbiome diversity, each measure of degree centrality, and for each species. Each of these full models was reduced using backwards stepwise model selection by AIC. Finally, we corrected p-values for false-discovery-rate with a Benjamini-Hochberg correction.

## RESULTS

We captured 40 mice and 91 voles. Of these, we had enough data to build the social network for 27 mice (20 males, 7 females) and 48 voles (26 males, 21 females, 1 unknown sex) and, of these, we obtained viable microbiome samples from 18 mice (17 males, 1 female) and 34 voles (24 males, 10 females). The social network constructed from both species, as well as those of each species, were more structured than random (Figure 1, S3). We tested for an effect of species and sex on degree centrality (linear model with species and sex as fixed effects) to assess how an over-representation of male mice might affect the interpretation of our results. Degree centrality of mice was lower than that of voles and this was driven by a lower degree centrality in female mice compared to male mice (species (vole) × sex (male): t=−3.43, P<0.001, R^2^_adj_=0.21; Table S3).

We used the scores of the first three axes of a PCA (Figure S5 and S6) on the habitat community data measured at each trapping location as habitat variables when testing our hypotheses about social proximity and microbiome structure and diversity. These axes accounted for 24.1%, 11.2%, and 7.4% of the variance in habitat data. Axis 1 (Habitat-PC1) represented a gradient going from open, coniferous forest with low fruiting shrubs (e.g. blueberry; *Vaccinium myrtilloides*) to mixed forest with coarse woody debris. Axis 2 (Habitat-PC2) represented a gradient going from coniferous forest with low understory made up of mosses, starflower (*Trientalis borealis*) and bunchberry (*Cornus canadensis*), to poplar-dominated deciduous forest with an understory of beaked hazel (*Corylus cornuta*) and red osier dogwood (*Cornus stolonifera*). Finally, axis 3 (Habitat-PC3) represented a gradient going from poplar-dominated deciduous forest with soils relatively high in moisture and pH, to mixed forest with low-moisture, low-pH, litter-covered soil.

The GM community dissimilarity of mice and voles were correlated with the social network (Procrustes analyses; mice: m_12_^2^ =0.32, t=0.84, P<0.001; voles: m_12_^2^ =0. 47, t=0.74, P=0.01) and habitat dissimilarity (mice: m_12_^2^ =0.48, t=0.64, P<0.001; voles: m_12_^2^ =0.63, t=0.63, P=0.02). For the full dual-species network, GM community dissimilarity correlated with the social network (m_12_^2^ =0.68, t=0.56, P<0.001), but not with habitat dissimilarity (m_12_^2^ =0.86, t=0.37, P=0.15). Similarly, for mouse-vole pairs, GM distance remained correlated with social distance (m_12_^2^ =0.70, t=0.55, P=0.03), but not with habitat distance (m_12_^2^ =0.84, t=0.40, P=0.22).

When analysing the effect of social distance on GM composition accounting for habitat by RDA, Mouse GM composition was positively correlated with Habitat-PC2 (F=2.05, P=0.001) and Habitat-PC3 (F=1.33, P=0.016; Figure 2). The adjusted cumulative R-squared (R^2^_adjcum_) for these variables was 0.077. When we repeated the modelling procedure with habitat removed, GM correlated with Social-PCo1 (F=1.85, P=0.001) and Social-PCo2 (F=1.43, P=0.003) and both explanatory variables had a R^2^_adjcum_ of 0.072 (Figure 3). Finally, when we repeated the above procedure while conditioning on habitat, GM was no longer correlated with the social network.

**Figure 3.**
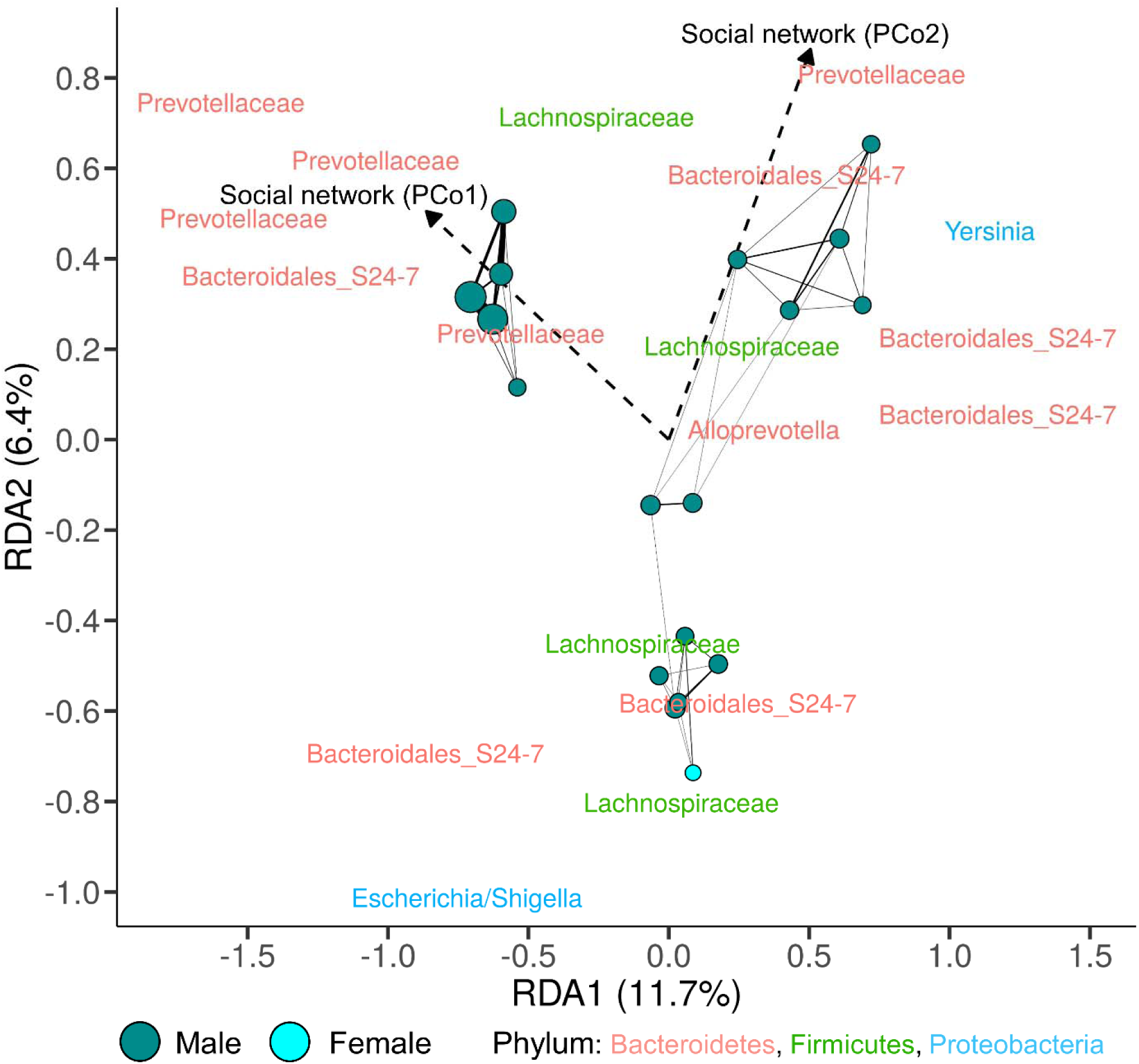
Biplot of RDA of the association between GM composition and social proximity in *P. maniculatus*. The model was selected from a full model that included only social network variables as explanatory factors. See Figure 2 for a description of the plot.

Vole GM composition correlated with Habitat-PC2 (F=1.48, P=0.007; Figure 4), Social-PCo1 (F=1.52, P=0.008) and sex (F=1.35, P=0.03) and these variables had a R^2^_adjcum_ of 0.04. When we removed habitat from the full model, GM was correlated with Social=PCo1(F=1.48, P=0.005) and Social-PCo2 (F=1.40, P=0.02), and was also correlated with sex (F=1.34, P=0.04; R^2^_adjcum_ =0.04; Figure 5). When we conditioned on habitat and sex, GM composition remained correlated with Social-PCo1(F=1.66, P=0.003; R^2^_adjcum_ =0.06).

**Figure 4.**
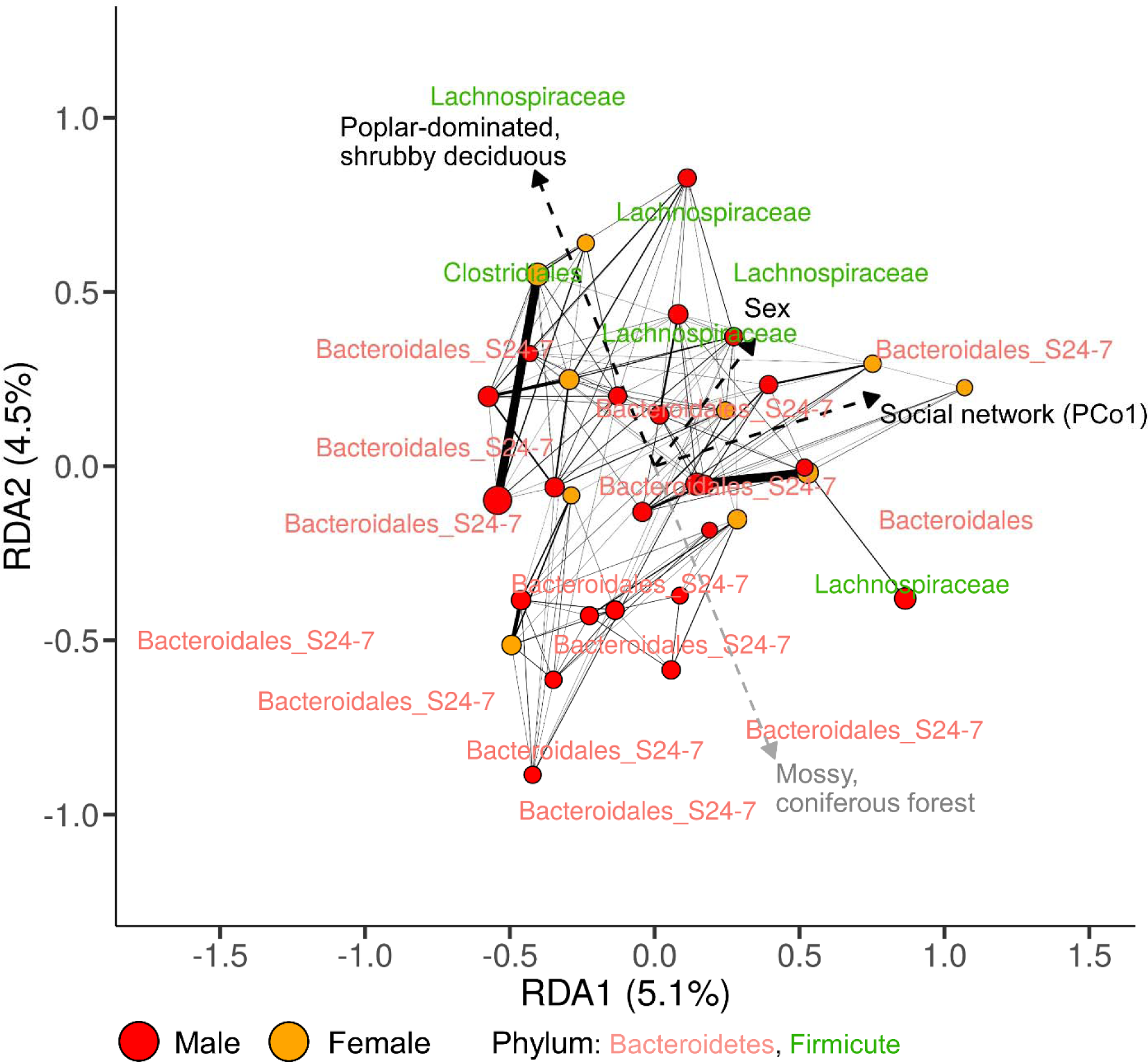
Biplot of RDA of the association between GM composition and social network, habitat and sex in *M. gapperi*. The RDA model was selected from a full model that included social network, habitat, and sex as explanatory factors. Variables correlated with GMcomposition and the direction and strength of the correlations are indicated by arrows. Habitat variables used were scores of a PCA of the habitat data and are represented above by decriptions of variation along the PCA axes. Social variables used were scores of a PCoA on social distance. Individuals with the strongest social associations (above the third quartile of social distances) are connected by lines, line width is proportional to the strength of association, and point size is proportional to degree centrality in the vole social network. The 10 most strongly loaded bacterial ASVs along each axis (identified to the finest-level taxonomic annotation for that ASV) are plotted (duplicate ASVs were removed).

**Figure 5.**
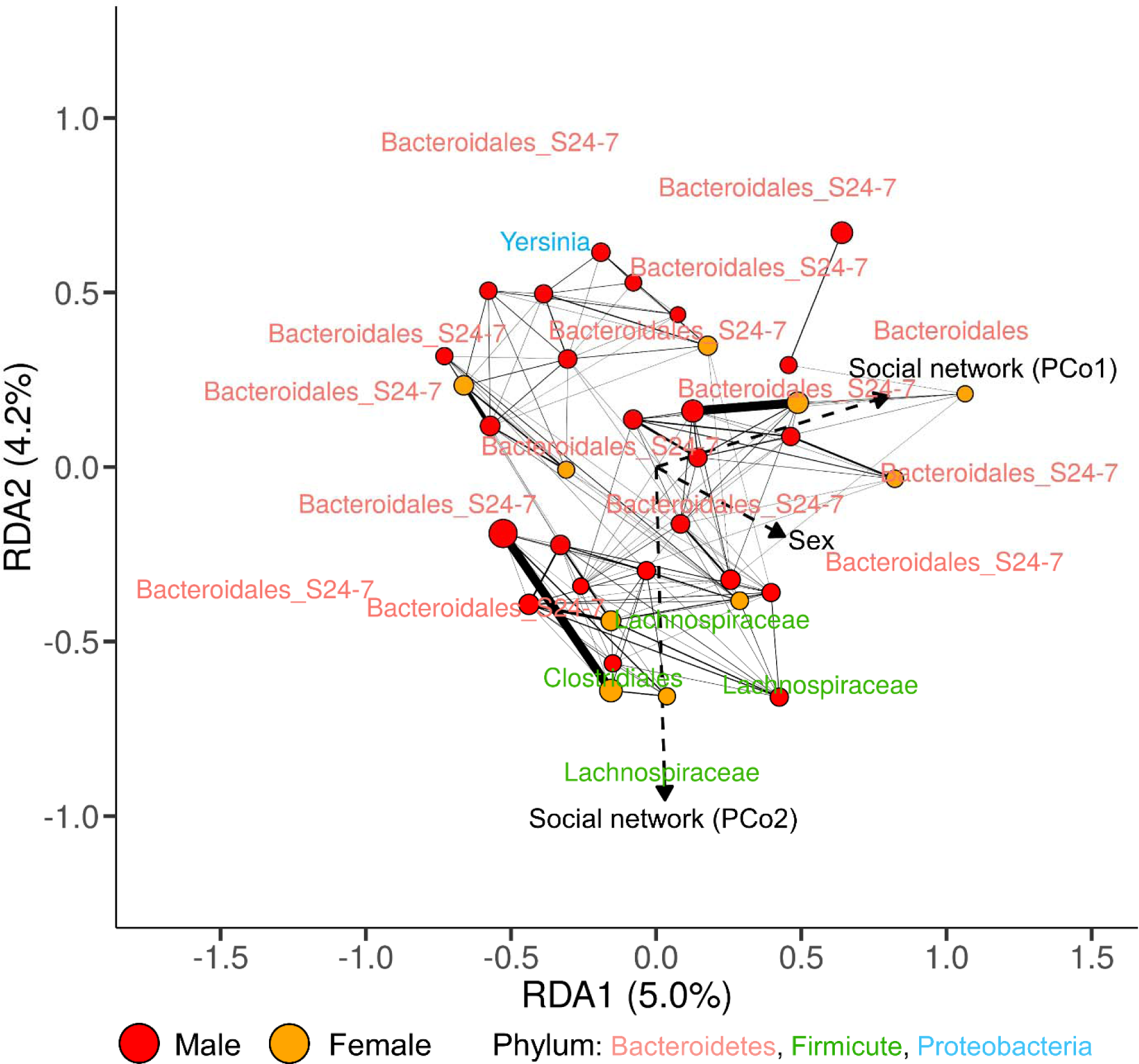
Biplot of RDA of the association between microbiome composition and social proximity and sex in *M. gapperi*. The model was selected from a full model that included social network and sex as explanatory factors. See Figure 4 for a description of the plot.

When we performed a combined analysis including mice and voles, conditioning on sex and species, GM composition was only correlated with Habitat-PC3 (F=2.43, P=0.005, R^2^=0.03). GM composition did not correlate with social proximity when we excluded habitat from the analyses since no variables were retained in the model during the stepwise model selection procedure.

We tested if mice GM diversity increased with increasing degree centrality in the mouse social network. We only found an association between GM Shannon’s diversity and Habitat-PC2 and between GM richness and both Habitat-PC1 and Habitat-PC2 (Table 1). When we regressed mouse GM diversity on habitat and degree centrality obtained from the dual-species network, GM Shannon’s diversity and GM richness both correlated with Habitat-PC1 and Habitat-PC2, but GM Shannon’s diversity also correlated with social degree centrality (Table 1, Table S1). Finally, when we tested for a relationship between GM diversity and social degree centrality of each mouse with respect to the vole network, GM Shannon’s diversity and richness correlated with Habitat-PC1 and social degree centrality (Table 1, Table S1). Additionally, using social degree centrality with respect to the vole network resulted in models that explained four times as much of the mouse GM Shannon’s diversity and twice as much of mouse GM richness as including degree centrality with respect to the mouse network (Table 1, Table S1). Simpsons’ diversity of the mouse GM did not correlate with habitat or any measure of social degree centrality (Table 1, Table S1).

**Table 1.**
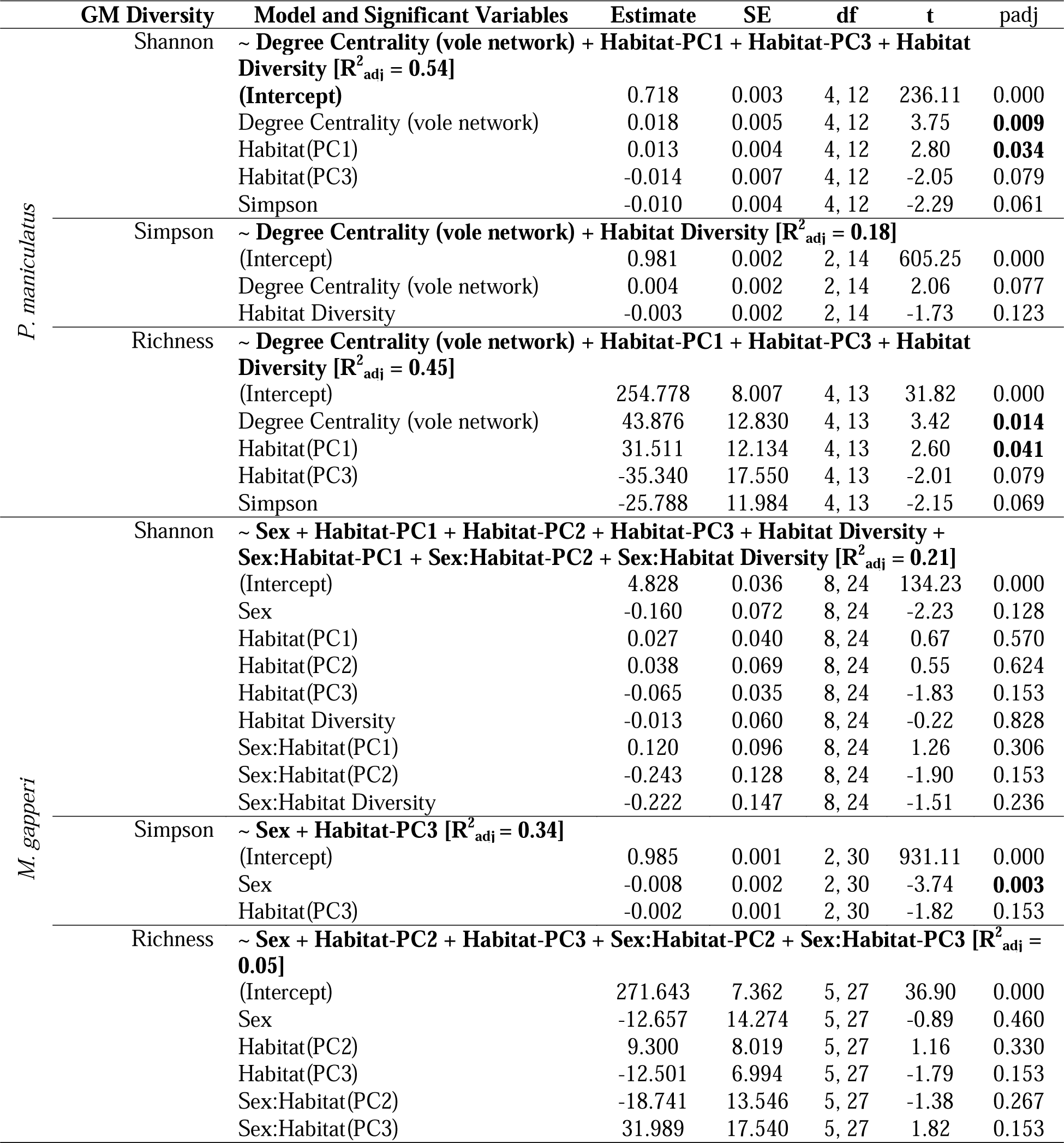
Separate linear regression analyses were conducted to test for a relationship between three measures of GM diversity (response; Shannon’s, Simpson, and Richness) and host social degree centrality (predictor) in mice and voles. To compare within- and cross-species interactive effects on GM diversity, for each species and each diversity metric, we conducted separate analyses with degree centrality of each individual with respect to either: 1) its species’ social network, 2) the dual-species network, and 3) the other species’ network. We accounted for habitat composition and diversity in analyses. Final models were derived by AIC-based stepwise selection and the best model for each measure of GM diversity is presented here (see table S1 for results of the final models from analyses with each measure of degree centrality).

In contrast to mice, the only association with vole GM diversity we found was between GM Simpson’s diversity and sex, with males having higher diversity than females (Table 1, Table S2).

## DISCUSSION

We hypothesized that similarity in GM composition between hosts, within and across host species, would correlate with their social distance as measured by their probability of encounter. We found that, in mice, social position and habitat were related, and that habitat was a slightly better predictor of GM composition. In contrast, in voles, social position best explained part of their GM composition and a second component of vole GM varied with both habitat and social position but was best explained by habitat. Finally, we found some evidence for interspecific interactions driving GM composition. Our second hypothesis was that GM diversity would vary with the number of neighbours in the social network, both within and cross-species. This hypothesis was only supported for the number of vole neighbours on the GM of mice.

The confounding effects of social position and habitat in mice made it impossible to confirm a role for social distance in shaping the mouse GM composition. This is likely a result of the method (Wanelik and Farine, 2019) we used to generate the social network. We used data on spatial occupation to estimate social connections and habitat is also spatially structured, rendering both factors highly related and difficult to dissociate. While obtaining observations of true contacts may provide a more accurate estimate of the social network, these may be rare (especially for less social individuals) and difficult to capture. Our method allowed estimation of both strong and weak social connections. Additionally, our hypotheses were based on a host-as-patch paradigm which does not preclude indirect transmission between hosts via the environment and thus probability of transmission should still depend on host isolation. Separating social distance and habitat effects on GM in mice will require sampling in areas where habitat is homogeneous.

Given that our measure of social distance among individuals depended on their movement, the effect of social distance we observed may be a result of individual differences in spatial behaviours such as exploration. GM composition in wild deer mice has been shown to directly influence host exploration in an open field with consequential effects on home range size (Jameson et al., 2020). We found that Bacteroidales-S24-7 (phylum Bacteroidetes) and Lachnospiraceae (phylum Firmicutes) strongly associated with position in the social network in both mice and voles (see figure 3 and 5), and these taxa have been shown to associate with open-field exploration in wild deer mice (Jameson et al., 2020). Fast explorers should have larger home ranges and have a greater probability of encountering more conspecifics and heterospecifics, and experience a greater diversity of habitats. All these features should impact their GM. The GM could also directly influence social behaviour such that a GM may promote host cohesive behaviour, rendering its hosts more likely to associate with each other, which could consequently facilitate its transmission among the hosts. Some members of Bacteroidales-S24-7 are also associated with social behaviour, with their abundance linked to social and cognitive impairment and depression in a mouse model of autism (Golubeva et al., 2017). That this bacterial group was strongly associated with social distance in both mice and voles, suggests it could be associated with social behaviour in these species and that it may be transmissible between hosts.

Transmission of bacteria between hosts could also occur through indirect routes of transmission. For example, members of the Enterobacteriaceae (phylum Proteobacteria), *Escherischia/Shigella* in mice and *Yersinia* in mice and voles, were associated with variation in social position and are transmitted via the fecal-oral route. However, these bacteria are most commonly dispersed by water (Percival and Williams, 2014) and, indeed, we found that this group associated most with humid habitats. These habitats may have facilitated transmission of these taxa among individuals. This could explain the confounded habitat and social effects we observed on the GM. Behaviours such as scent marking may also facilitate transmission of these and other bacterial species. Scent marking was proposed as a driver of socially transmitted *E. coli* in Verreaux’s sifakas (*Propithecus verreauxi*) (Springer et al., 2016). In rodents, urine and feces-derived chemical cues are used to communicate identity and dominance, mark territories, navigate, select and attract mates, and influence social behaviour and reproductive physiology of conspecifics (Johnston, 2003; Ma et al., 1999). Contact with these bacterial sources offers an indirect mechanism for the social transmission of bacteria. Additionally, mice and voles are coprophagic (Cranford and Johnson, 1989), which could also drive bacterial transmission in these species, yet the extent to which they consume the feces of other individuals or species is unclear.

Finally, when we considered the interactions between both species on GM composition, Procrustes analysis supported our hypothesis that individuals of different species with more frequent interactions share greater similarity in their GM composition. While we did not observe this with the RDA, Procrustes targeted mouse-vole interactions which made it more suited to testing our hypothesis. Additionally, while we attempted to correct for species differences in the RDA, transmission between species may be confounded with species differences in GM. This is because taxa transmitted between species may also be common in one species but rare in the other and thus be removed when correcting for species differences. In any case, our results are consistent with bacterial dispersal within and between species following metacommunity theory, though the effect of potential interactions between mice and voles on their GM is likely weak.

We predicted that the number of potential encounters (i.e. degree centrality) would promote transmission of bacteria from different communities and result in greater GM diversity. This was not the case for either species when we examined degree centrality measured within each species’ conspecific network. Surprisingly, degree centrality of mice with respect to voles, and to a lesser degree, that of mice within the dual-species network, were strong predictors of mouse GM diversity. Though the lower density of mice may have limited our ability to detect a correlation between intraspecific degree centrality and GM diversity, these results provide further evidence for an influence of bacterial transmission across different host species on the GM of these animals. As far as we know, this has not yet been shown in wild species. GM diversity measures associated with degree centrality were sensitive to rare taxa suggesting that cross-species transfers mainly involved rare taxa. As such, bacteria obtained by mice from voles are more likely to be transient or may not share a commensal relationship with their new host.

We observed a robust association between GM composition and social proximity in voles. In mice, social proximity was correlated with both habitat and GM composition. Our measures of habitat may not account for some components of diet (e.g. invertebrates), and diet could be a stronger driver of spatial variation in GM than host-host transmission (David et al., 2014; Ley et al., 2006; Muegge et al., 2011). While some studies have found no evidence of deer mice forming long-lasting social groups with kin (Dewsbury, 1990), other studies suggest they can form family groups (Savidge, 1974) and demonstrate female-biased philopatry (Teferi, 1993). Thus, another factor that may be confounded with social distance is genotype and bacteria may also be transferred from parent to offspring and among kin in the natal nest. However, Schmidt et al. (2019) observed a weak effect of genotype on within-population GM composition of deer mice, and our hypotheses focus on host-host dispersal as a driver of GM composition which does not preclude dispersal among kin. We also found fundamental differences in the network structure of mice and voles based on sex with female mice less central than male mice and found no such difference in voles. This difference in network structures suggests that associations between GM and frequency of encounters in mice may differ between sexes. Expression of aggressive behaviours differs between male and female deer mice but such differences are less evident in red-backed voles (Dewsbury, 1981; Dracup et al., 2016; McGuire, 1997; Mihok, 1981). These behavioural differences could lead to sex and species-specific differences in bacterial transmission dynamics.

Our results demonstrate, besides the mounting evidence from highly social species such as non-human primates (Moeller et al., 2016; Raulo et al., 2018; Tung et al., 2015) that the GM composition and diversity of less social species follow a metacommunity theory framework. Of exceptional interest is the finding that the GM of a species is likely influenced by the GM of surrounding heterospecifics. The GM may therefore play a role in regulating host communities, either through dispersal of beneficial or pathogenic bacteria. Despite this, probability of contact among hosts explained a relatively small proportion of the GM in mice and voles. While metacommunity theory offers a convenient starting point to understand how host behaviourally-driven microbial dispersal shapes the GM, it currently does not account for the host’s capacity to directly influence its GM and the GM’s capacity to influence host health and behaviour. Integrating these host-GM feedbacks in current metacommunity theory should greatly improve its ability to explain GM composition within and across host species.

## ACKNOWLEDGMENTS

Animal care and experimental procedures were performed following protocols approved by the Comité institutionnel de protection des animaux (CIPA #917). We thank S. Zhao, C. Pelletier, R. Pedneault for their assistance and dedication in the field and K. and B. Hall for their logistical support in the field. C. Negre performed the DNA extractions and assisted with analyses. We thank D. Farine and K. Wanelik for their suggestions and support with social network analyses. J. Jameson received an NSERC Alexander Graham Bell Canada Graduate Scholarship, and S. Zhao and C. Pelletier received NSERC-USRA fellowship. This research was funded by Natural Sciences and Engineering Research Council of Canada (NSERC) Discovery grants (Réale and Kembel), a Fonds de recherche du Québec – Nature et technologies (FRQNT) Team Grant (Réale and Kembel), a Canada Research Chair (Kembel), an American Society of Mammalogists Grant-in-Aid of Research (Jameson) and an Animal Behavior Society Student Research Grant (Jameson).

## SUPPLEMENTAL MATERIAL

**Figure S1.**
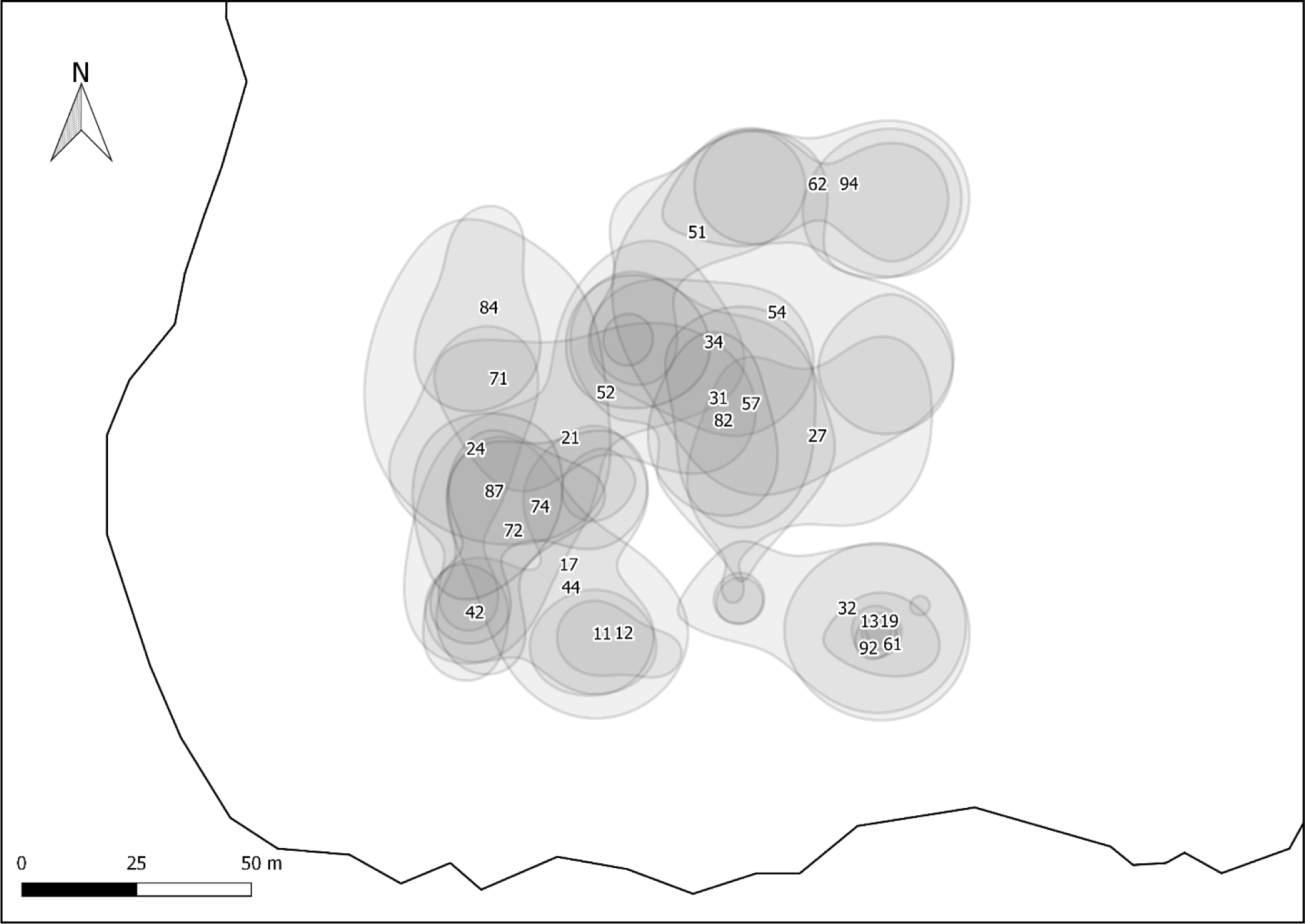
Home ranges (50% kernel, package adehabitatHR) of *P. maniculatus* based on live captures and PIT tag detections at RFID antennae. Numbers represent the individual’s IDs.

**Figure S2.**
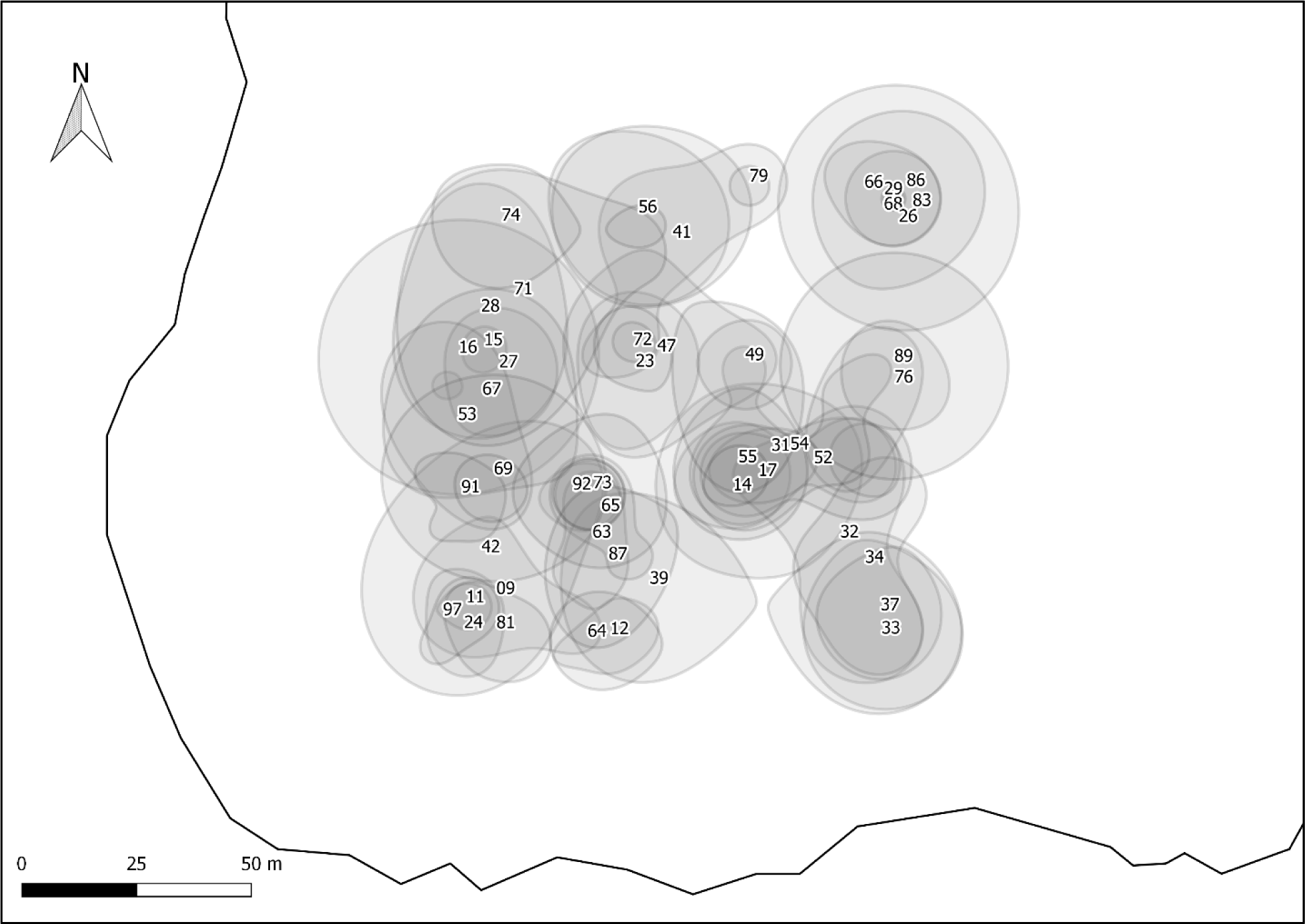
Home ranges (50% kernel, package adehabitatHR) of voles based on live captures and PIT tag detections at RFID antennae. Numbers represent the individual’s IDs.

**Figure S3.**
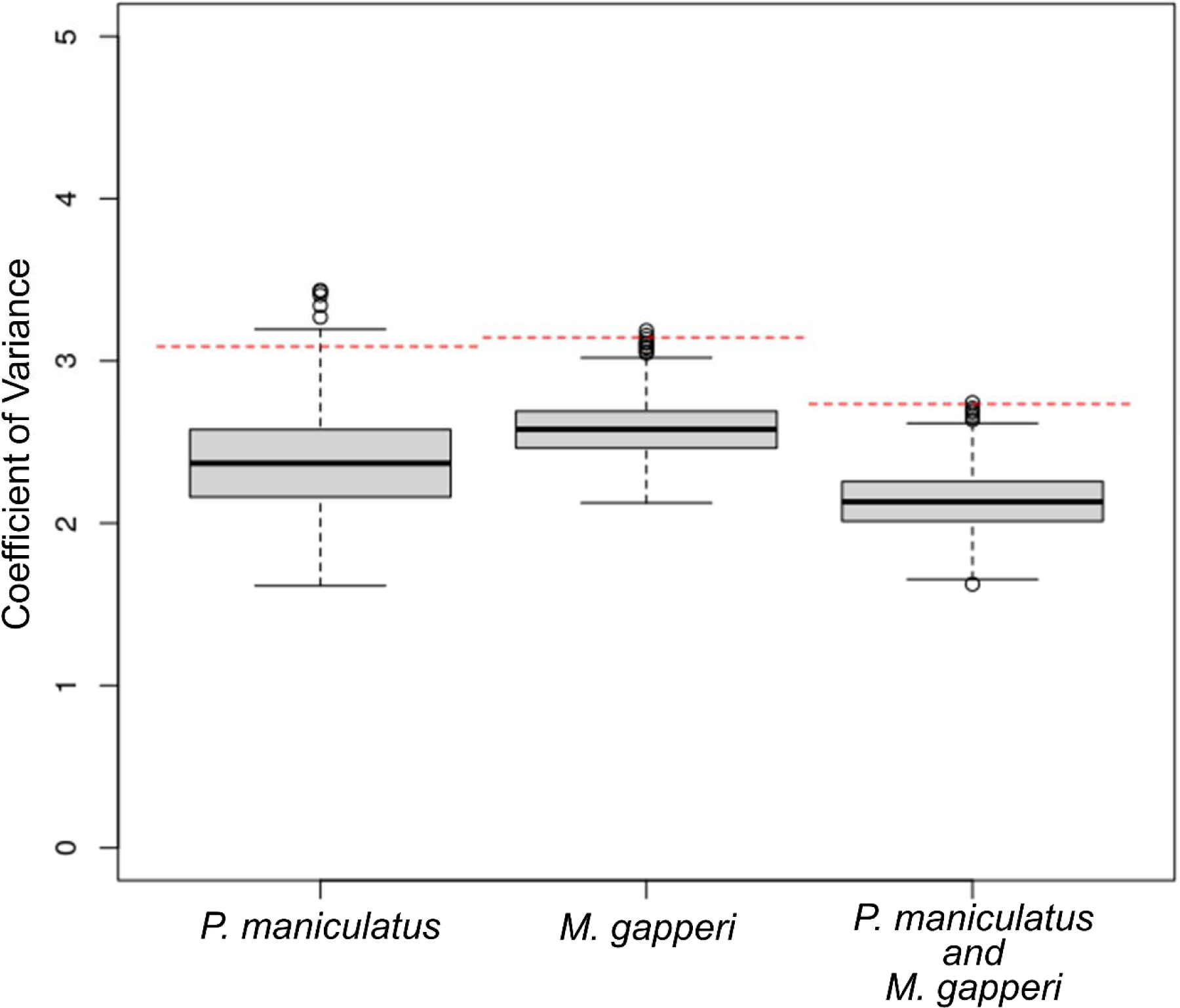
Boxplots of results from permutation tests to determine if social networks constructed for *P. maniculatus*, *M. gapperi*, and both species together, are more structured than random. Boxplots are derived from the coefficient of variance (y axis) for 1000 random permutations of the social network. The dotted red lines represent the coefficients of variance of the observed social networks (CVobs).

**Figure S4.**
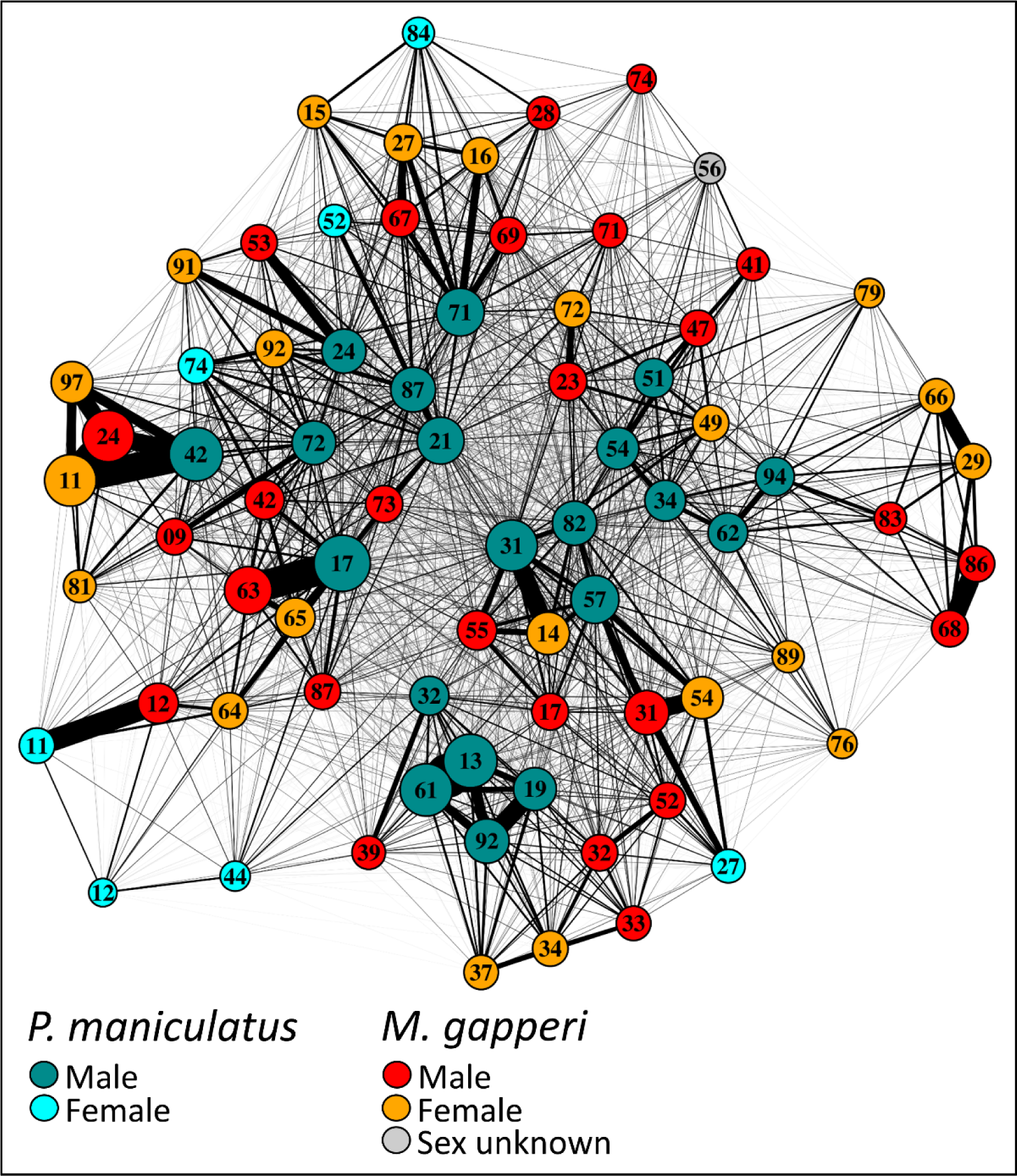
Combined social network of mice and voles. Vertex size is proportional to the degree centrality of the individual in the social network. Edge width is proportional to the strength of association. Numbers identify individuals.

**Figure S5.**
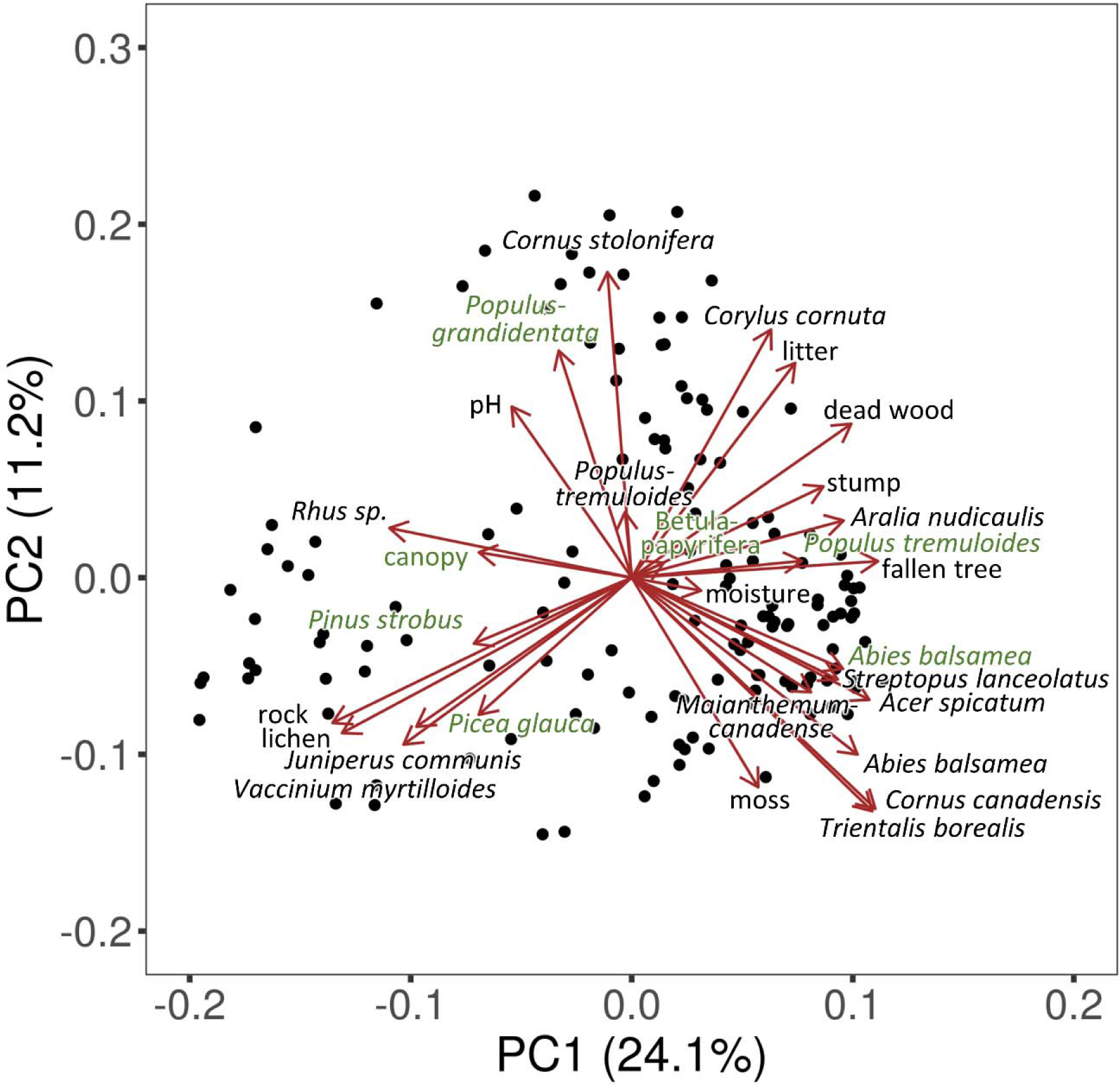
Biplot of the first and second axes of a principal components analysis of habitat community data collected at each trapping location (points in the figure). Species comprising the overstory stratum are indicated in green.

**Figure S6.**
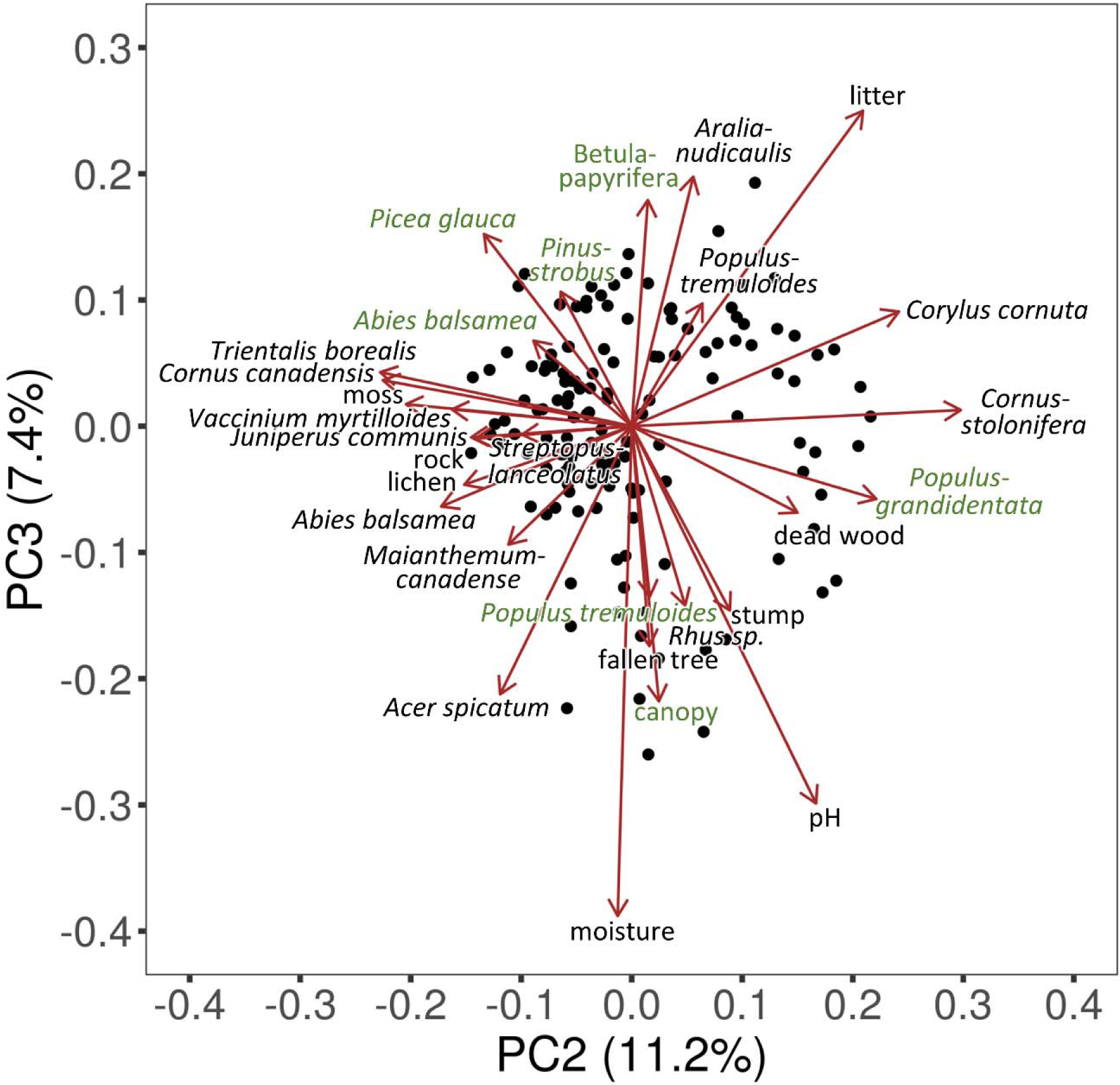
Biplot of the second and third axes of a principal components analysis of habitat community data collected at each trapping location (points in the figure). Species comprising the overstory stratum are indicated in green.

**Table S1.** Separate linear regression analyses were conducted to test for a relationship between three measures of GM diversity (response; Shannon’s, Simpson, and Richness) and host social degree centrality (predictor) in mice. To compare within- and cross-species interactive effects on GM diversity, for each species and for each measure of diversity, we conducted separate analyses including one of three measures of degree centrality: degree centrality of each individual with respect to its species’ social network, degree centrality with respect to the dual-species network, and degree centrality of each individual with respect to the other species’ network. We accounted for habitat composition and diversity in analyses. In each analysis, a model was derived using stepwise model selection (comparing AIC values) on the following full model: diversity ~ degree centrality + Habitat-PC1 + Habitat-PC2 + Habitat-PC3 + Habitat Diversity.

| GM Diversity | Model and Variables | Estimate | SE | df | t | padj | LCI | UCI |
| --- | --- | --- | --- | --- | --- | --- | --- | --- |
| Shannon | ~ Habitat-PC1 + Habitat-PC2 + Habitat-PC3 [$R^2_{adj} = 0.17$] | | | | | | | |
|  | (Intercept) | 0.718 | 0.004 | 3, 13 | 175.54 | 0.000 | 0.709 | 0.727 |
|  | Habitat(PC1) | 0.020 | 0.009 | 3, 13 | 2.26 | 0.061 | 0.001 | 0.039 |
|  | Habitat(PC2) | 0.028 | 0.011 | 3, 13 | 2.47 | **0.046** | 0.004 | 0.053 |
|  | Habitat(PC3) | 0.010 | 0.007 | 3, 13 | 1.43 | 0.186 | -0.005 | 0.026 |
| | ~ Degree Centrality(mouse/vole network) + Habitat-PC1 + Habitat-PC2 [$R^2_{adj} = 0.38$] | | | | | | | |
|  | (Intercept) | 0.718 | 0.004 | 3, 13 | 202.42 | 0.000 | 0.710 | 0.726 |
|  | Degree Centrality(mouse/vole network) | 0.011 | 0.004 | 3, 13 | 2.65 | **0.040** | 0.002 | 0.020 |
|  | Habitat(PC1) | 0.022 | 0.008 | 3, 13 | 2.91 | **0.029** | 0.006 | 0.039 |
|  | Habitat(PC2) | 0.018 | 0.007 | 3, 13 | 2.48 | **0.046** | 0.002 | 0.034 |
| | ~ Degree Centrality(vole network) + Habitat-PC1 + Habitat-PC3 + Habitat Diversity [$R^2_{adj} = 0.54$] | | | | | | | |
|  | (Intercept) | 0.718 | 0.003 | 4, 12 | 236.11 | 0.000 | 0.711 | 0.725 |
|  | Degree Centrality(vole network) | 0.018 | 0.005 | 4, 12 | 3.75 | **0.009** | 0.008 | 0.029 |
|  | Habitat(PC1) | 0.013 | 0.004 | 4, 12 | 2.80 | **0.034** | 0.003 | 0.022 |
|  | Habitat(PC3) | -0.014 | 0.007 | 4, 12 | -2.05 | 0.079 | -0.028 | 0.001 |
|  | simpson | -0.010 | 0.004 | 4, 12 | -2.29 | 0.061 | -0.020 | -0.001 |
| Simpson | No suitable model with Degree Centrality(mouse network) |  |  |  |  |  |  |  |
|  | (Intercept) | 0.981 | 0.002 | 0, 16 | 548.30 | 0.000 | 0.977 | 0.985 |
| | ~ Degree Centrality(mouse/vole network) + Habitat-PC1 + Habitat Diversity [$R^2_{adj} = 0.09$] | | | | | | | |
|  | (Intercept) | 0.981 | 0.002 | 3, 13 | 575.42 | 0.000 | 0.977 | 0.985 |
|  | Degree Centrality(mouse/vole network) | 0.003 | 0.002 | 3, 13 | 1.72 | 0.123 | -0.001 | 0.008 |
|  | Habitat(PC1) | 0.003 | 0.002 | 3, 13 | 1.32 | 0.211 | -0.002 | 0.008 |
|  | simpson | -0.003 | 0.002 | 3, 13 | -1.32 | 0.211 | -0.008 | 0.002 |
| | ~ Degree Centrality(vole network) + Habitat Diversity [$R^2_{adj} = 0.18$] | | | | | | | |
|  | (Intercept) | 0.981 | 0.002 | 2, 14 | 605.25 | 0.000 | 0.978 | 0.985 |
|  | Degree Centrality(vole network) | 0.004 | 0.002 | 2, 14 | 2.06 | 0.077 | 0.000 | 0.007 |
|  | Habitat Diversity | -0.003 | 0.002 | 2, 14 | -1.73 | 0.123 | -0.007 | 0.001 |
| Richness | ~ Habitat-PC1 + Habitat-PC2 + Habitat-PC3 [$R^2_{adj} = 0.25$] | | | | | | | |
|  | (Intercept) | 254.778 | 9.612 | 3, 14 | 26.51 | 0.000 | 234.162 | 275.393 |
|  | Habitat(PC1) | 55.201 | 21.555 | 3, 14 | 2.56 | **0.041** | 8.970 | 101.432 |
|  | Habitat(PC2) | 80.723 | 27.795 | 3, 14 | 2.90 | **0.029** | 21.108 | 140.338 |
|  | Habitat(PC3) | 26.559 | 16.805 | 3, 14 | 1.58 | 0.150 | -9.484 | 62.602 |
| | ~ Degree Centrality(mouse/vole network) + Habitat-PC1 + Habitat-PC2 [$R^2_{adj} = 0.34$] | | | | | | | |
|  | (Intercept) | 254.778 | 9.043 | 3, 14 | 28.17 | 0.000 | 235.382 | 274.173 |
|  | Degree Centrality(mouse/vole network) | 22.890 | 10.629 | 3, 14 | 2.15 | 0.069 | 0.094 | 45.687 |
|  | Habitat(PC1) | 58.741 | 20.396 | 3, 14 | 2.88 | **0.029** | 14.997 | 102.486 |
|  | Habitat(PC2) | 53.689 | 19.510 | 3, 14 | 2.75 | **0.034** | 11.844 | 95.534 |
| | ~ Degree Centrality(vole network) + Habitat-PC1 + Habitat-PC3 + Habitat Diversity [$R^2_{adj} = 0.45$] | | | | | | | |
|  | (Intercept) | 254.778 | 8.007 | 4, 13 | 31.82 | 0.000 | 237.480 | 272.076 |
|  | Degree Centrality(vole network) | 43.876 | 12.830 | 4, 13 | 3.42 | **0.014** | 16.157 | 71.594 |
|  | Habitat(PC1) | 31.511 | 12.134 | 4, 13 | 2.60 | **0.041** | 5.297 | 57.725 |
|  | Habitat(PC3) | -35.340 | 17.550 | 4, 13 | -2.01 | 0.079 | -73.255 | 2.576 |
|  | simpson | -25.788 | 11.984 | 4, 13 | -2.15 | 0.069 | -51.677 | 0.102 |

**Table S2.**
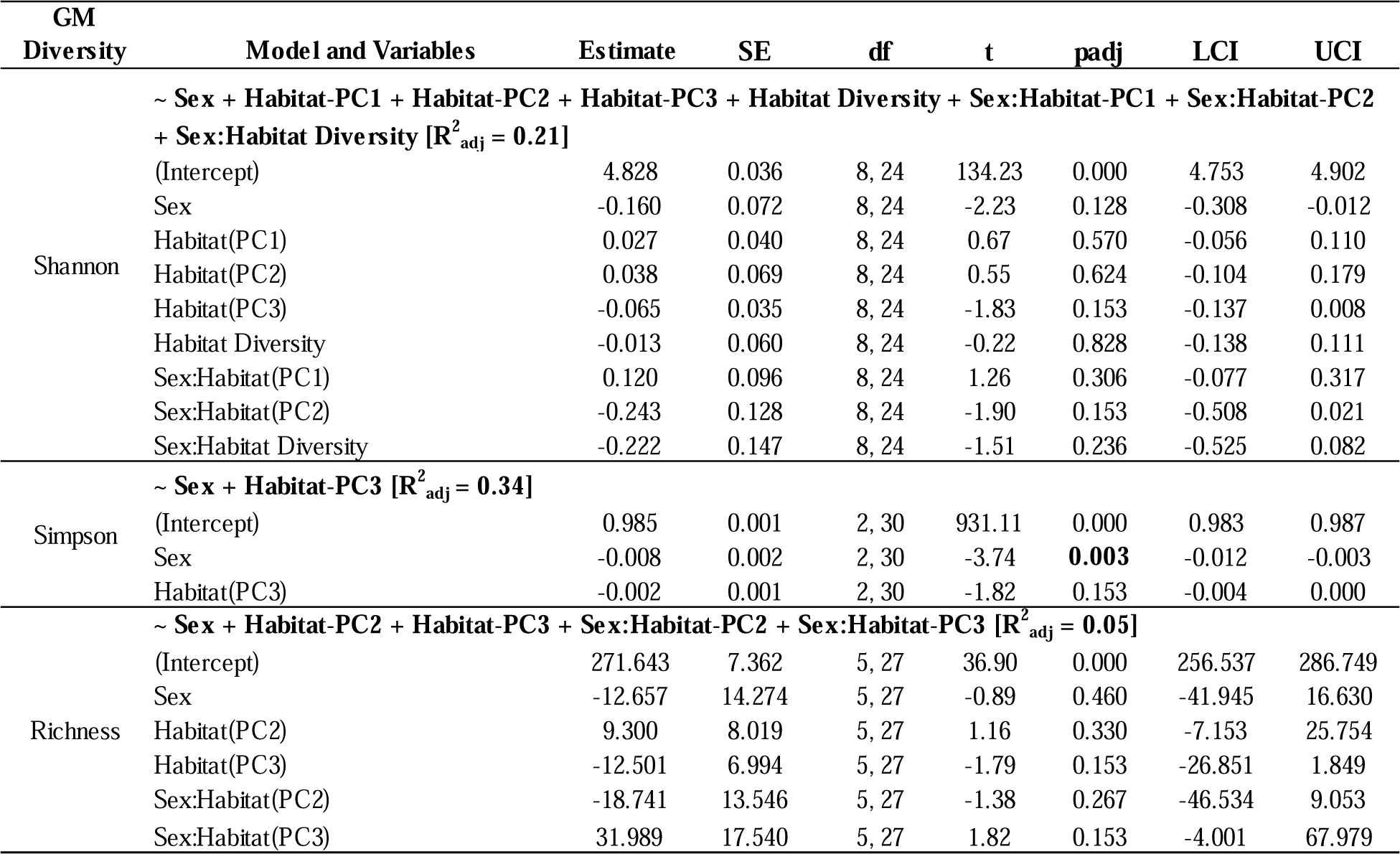
Separate linear regression analyses were conducted to test for a relationship between three measures of GM diversity (response; Shannon’s, Simpson, and Richness) and host social degree centrality (predictor) in voles. To compare within- and cross-species interactive effects on GM diversity, for each species and for each measure of diversity, we conducted separate analyses including one of three measures of degree centrality: degree centrality of each individual with respect to its own species’ social network, degree centrality with respect to the dual-species network, and degree centrality of each individual with respect to the other species’ network. We accounted for habitat composition and diversity in analyses. In each analysis, a model was derived using stepwise model selection (comparing AIC values) on the following full model: diversity ~ degree centrality + Habitat-PC1 + Habitat-PC2 + Habitat-PC3 + Habitat Diversity. In voles, final models did not contain degree centrality and therefore, the same model was obtained within each measure of GM diversity regardless of the measure of degree centrality used.

**Table S3.** Results from a linear regression model testing for an effect of species (Mouse and Vole) and sex (Male and Female) on degree centrality of individuals in their respective species’ social network.

| Variable | Estimate | SE | t | p | LCI | UCI |
| --- | --- | --- | --- | --- | --- | --- |
| (Intercept) | -1.45 | 0.15 | -9.41 | <0.001 | -1.75 | -1.14 |
| Species(V) | 0.65 | 0.18 | 3.67 | <0.001 | 0.30 | 1.01 |
| Sex(M) | 0.82 | 0.18 | 4.56 | <0.001 | 0.46 | 1.17 |
| Species(V):Sex(M) | -0.74 | 0.22 | -3.44 | 0.001 | -1.17 | -0.31 |

### SEQUENCE DATA QUALITY CONTROL METHODS

Following quality control, filtering, and removal of chimeras, we identified reads to amplicon sequence variants (ASVs) with package DADA2 version 1.10 (Callahan et al., 2016). We used default settings for all DADA2 analysis parameters specifying a forward read length of 250 and a reverse read length of 210 during read trimming. When merging forward and reverse reads, we specified a minimum overlap length of 10 and a maximum number of mismatches allowed in the overlap region of 10. We removed chimeras using the “consensus” method. This resulted in an average (±SD) of 18,878 ± 5,051 sequences per sample (minimum=4,775 sequences, maximum=39,164 sequences). We used the RDP Naïve Bayesian Classifier algorithm in DADA2 to assign taxonomy to ASVs according to the SILVA database (V128). We removed ASVs with <100 sequences and rarefied samples to 4 500 reads per sample, which was sufficient for rarefaction curves to reach a plateau without eliminating any samples for our dataset (Fig.S7).

**Figure S7.**
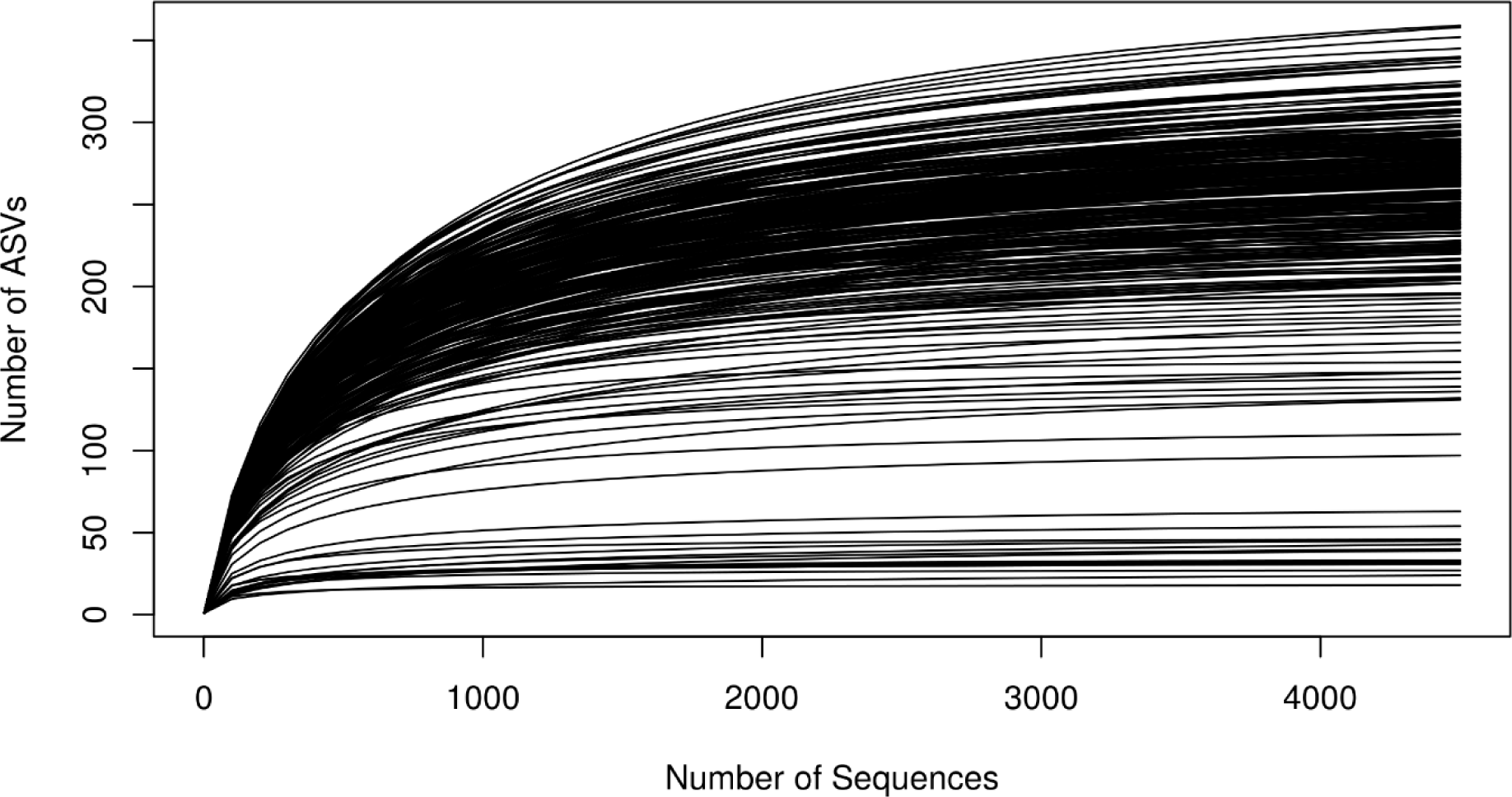
Rarefaction curves for all samples collected depicting the number of amplicon sequence variants (ASV) per number of sequences in each sample.

